# Gene Expression Signature and Molecular Mechanism of Redox Homeostasis in Colorectal Cancer

**DOI:** 10.1101/2020.01.28.920553

**Authors:** Mehran Piran, Maryam Darayee, Mehrdad Piran, Neda Sepahi, Amir Rahimi

## Abstract

Cellular redox homeostasis is the important tool for normal cell function and survival. Oxidants, reductants and antioxidants are the players to maintain cellular homeostasis balance. However, in some conditions like cancer, the concentration and activation of these players are disturbed. This study walks you through the molecular mechanism of redox homeostasis to describe how expression level of these players would help colorectal cancer (CRC) cells continue proliferation and survive in the hypoxic environment of tumor. We proposed that O_2_^-^ concentration is not detrimentally high in CRC cells since expression level of MnSOD didn’t change noticeably. We also suggested that High proliferative CRC cells obtain their energy by oxidation of H_2_S in or Electron transport chain (ETC) and keep the adequate concentration of H_2_S by diminishing the expression level of enzymes involved in sulfide oxidation pathway. Reduction in hydrogen sulfide oxidation results in a decrease in the level of GSH. Glutathione peroxidase enzyme requires GSH to convert H_2_O_2_ into oxygen and water. Therefore, Level of hydrogen peroxide stays high which leads to an increase in cell proliferation. Furthermore, we analyzed the expression level of transcription factors sensitive to redox messengers.

## Introduction

Redox homeostasis is the balance in concentration of cellular oxidants and reductants adequate to normal cell physiology [1]. This balance strongly relies on generation and eradication of redox messengers such as ROS (Reactive Oxygen Species), RNS (Reactive Nitrogen Species), H_2_S (Hydrogen Sulfide) and so on. These messengers are produced through enzymatic reactions or ETC and they are scavenged by antioxidants [2]. Oxidants grab electrons from proteins which leads to changes in proteins function and topological structure. They also attack Polyunsaturated fatty acids (PUFAs) in cell membranes causing irreparable damages [3].

Proteins and molecules which function as signaling messengers, transduce messages between cellular machines and determine how much ATP is consumed or produced in cells. Signaling messengers specify how cells respond to the environmental changes likewise cellular damages or changes in temperature or oxygen level. On the one hand ROS and RNS are the toxic byproducts of cellular respirations, on the other hand they have second messenger characteristics that control numerous cellular functions [4]. In other words, cells survival is not possible without redox molecules. A mild increase in the concentration of redox molecules leads to cell proliferation while a severe increment and long-lasting in the concentration of these molecules causes cell death. As a result, if redox homeostasis is disturbed, oxidative stress contributes to aberrant cell growth/death and disease [2]. This happens by changes in cells environment and impaired antioxidants generation which modulate the concentration of oxidants [5].

ROS includes radical species such as superoxide (O_2_^-^), hydroxyl radical (·OH) and non-radical species like Hydrogen peroxide (H_2_O_2_). RNS involves nitric oxide (NO) and Peroxynitrite (ONOO^-^) [2, 6]. Moreover, there are two types of antioxidants, enzymatic and non-enzymatic (small molecules). SODs (Superoxide dismutase), Glutathione peroxidase (GPx) and Catalase are examples of enzymatic antioxidants but Glutathione (GSH), Thioredoxin (Tnx) and Peroxiredoxin (Prx) are examples of small molecule antioxidants. One of the primary ROS molecules is O_2_^-^ made in the ETC by transferring electrons from complex one and Three to a dioxygen molecule or by NAD(P)H oxidases (NOX) activation [7–10]. Superoxide is then converted into hydrogen peroxide using Superoxide dismutase enzymes [9]. Among the three kinds of SODs, SOD1 (CuZnSOD) is present in the cytoplasm, mitochondrial inter-membrane space, nucleus, and lysosomes while SOD2 (MnSOD) and SOD3 (Ec-SOD) are present in Mitochondria and Extra Cellular Matrix (ECM) respectively [2]. Next, H_2_O_2_ is turned into water and oxygen molecules by GPx enzyme which oxidizes reduced Glutathione (GSH) into oxidized glutathione (GSSG) [11]. GSSG is converted into GSH by Glutathione reductase (GR/GSR) and in the meantime, an NADPH is turned into NADP^+^. GSH can enter the oxidation cycle again and NADP^+^ enters the glycolysis to be reduced into NADPH [12]. In the presence of Fe^+2^ and Fe^+3^, during Fenton and Haber-Weiss reactions, H_2_O_2_ can be converted into the harmful ·OH [2] which reflexes the essential role of SOD enzymes in Redox Homeostasis.

Redox homeostasis can control different stages of carcinogenesis including growth, metastasis, and angiogenesis [13, 14]. In addition, oxidants like ROS which are made during cell metabolism are vital for normal cell function and survival. As a case in point, DU-145 prostate cancer cells treated with low levels of H_2_O_2_ (100 nM to 1 mM) witness an increase in c-Fos expression which results in cell proliferation. However, if these cells are treated with 200 mM hydrogen peroxide, c-Fos expression is suppressed and DNA damage and cell cycle arrest lead to proliferation hindering. In addition to concentration of ROS molecules, exposure duration and location of redox molecules in different parts of a cell impact the apoptotic and antiapoptotic functions of these molecules [11].

In this study, we analyzed two transcriptome datasets from 108 Colorectal Cancer (CRC) patients from two studies with GEO accession ID GSE18105 (first dataset) and GSE41258 (second dataset). Each dataset contains three groups of samples including metastatic, primary and normal. A significant literature study was conducted to identify genes, proteins and antioxidants, along with a number of small-molecule redox messengers engaged in cellular redox homeostasis. To dissect redox mechanism of CRC progression, gene expression profiles of selected genes were extracted from two datasets and visualized with bar charts. Regarding the production and removal of redox messengers, the desired gene expressions were interpreted. Understanding the pathogenicity of these small messengers and their collaboration in promoting cancer, would help us not only introduce redox-related genes expression signature but also to identify specific CRC biomarkers for target therapy. Note that, a great number of genes were removed from the two datasets in the processing step. However, a few of them were important so we mentioned their expression results in Results and Discussion section.

## Methods

### Database and Literature Searching to Recognize Pertinent Experiments and Genes

Gene expression datasets were obtained from GEO repository which is one of the NCBI genomic databases (http://www.ncbi.nlm.nih.gov/geo/) holding high-throughput microarray and next-generation sequencing datasets. The keywords used to filter the search were human, colorectal/colon cancer, normal, primary, Benign, and metastasis. Two data sets with high-quality data each containing three groups of samples were retrieved. First dataset (GSE18105) contains 20 metastatic samples, 10 primary (benign) samples and 17 normal samples and second dataset (GSE41258) includes 31 metastatic samples, 22 primary samples and 8 normal samples. In the first dataset, metastatic samples were extracted from malignant colorectal tumors but in the second dataset, metastatic samples were obtained from liver metastasis tumors in CRC patients. primary samples were extracted from benign tumors (CRC stage I or II) and normal samples are normal colon tissue adjacent to colorectal tumors in both datasets. All the samples are presented in Supplementary file 1. In the next step, literature searching was practiced and a number of genes related to cellular redox homeostasis were selected.

### High-throughput Transcriptome Data Preprocessing

R software with version 3.6.1 was used to import and analyze data for each dataset separately. To make samples and datasets comparable, preprocessing step involving background correction and probe summarization was conducted using RMA method in Affy package [15]. Absent probesets were also identified using “mas5calls” function in this package. If half of the samples had absent values for a probeset, that probeset was regarded as absent and removed from the expression matrix. Outlier samples within each group in each dataset were removed based on PCA and hierarchical clustering approaches and box plot. After that, data were normalized using Quantile Normalization method. Many to Many problem [16] which is mapping multiple probesets to the same gene symbol was solved using nsFilter function in “genefilter” package [16]. Finally, expression data was plotted for genes engaged in the cellular redox homeostasis in two datasets.

## Results and Discussion

### Dataset Construction for Redox Homeostasis Genes

PCA was applied to datasets separately. Between three groups of samples, a number of them are away from their cluster so they were considered as outliers (Figure1A). To be more specific, a hierarchical clustering method introduced by Oldham MC [17] was used. Pearson correlation coefficients between samples were subtracted from one for measurement of the distances. Figure1B depicts the dendrogram for the normal samples. Figure1C normal samples are plotted based on their Number-SD scores. To get this number for each sample, the average of whole distances is subtracted from distances average in all samples then, results of these subtractions are normalized (divided) by the standard deviation of distance averages [17]. Samples with Number-SD less than negative two which were in a distance from their cluster set in the PCA plot were regarded as outliers. In addition, box plots for each dataset were plotted and samples with extreme IQRs were removed. Eleven outlier samples in the first dataset and twenty-one outlier samples in the second dataset were recognized. Supplementary file1 contains information about groups of samples and outliers. Finally, 85 genes engaged in cellular redox homeostasis were selected and 53 final genes were chosen based on their relationship with cancer progression. Bar plots for these gene expressions are illustrated in Supplementary file 1. Note that, the main interpretations were conducted on obtained results from first dataset however, results in the second dataset play a supportive role. Furthermore, some genes were found only in the first dataset.

**Figure1:**
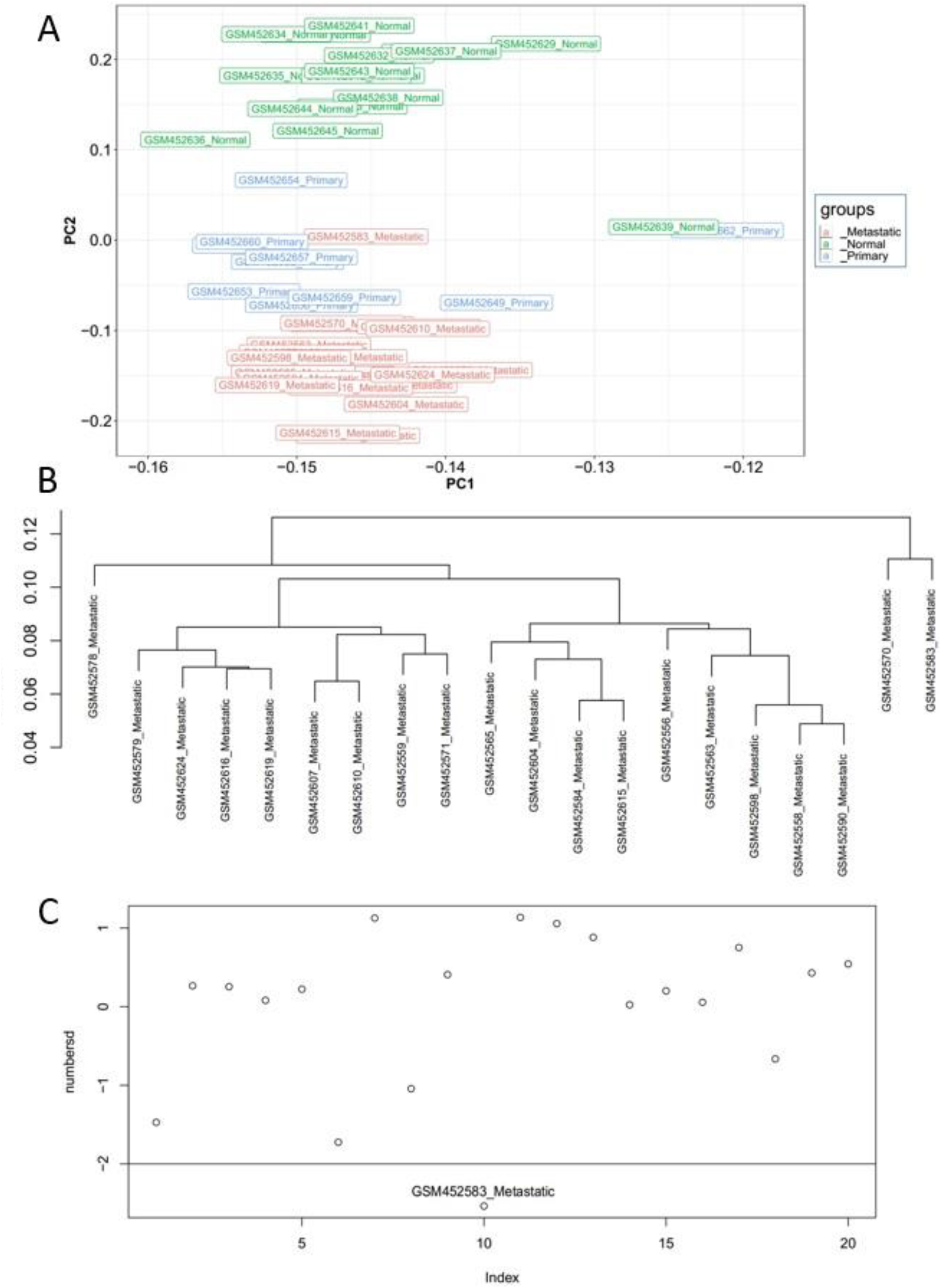
Quality control plots for the first dataset. A, illustrates the dispersion of samples based on eigenvecor1 (PC1) and eigenvector2 (PC2). B, is the dendrogram for normal group and C is the Number-SD plot. Sample GSM452639 in normal group is an outlier since in PCA, it is situated in the right-hand side, in dendrogram, it is in a distinct cluster and Its Number-SD is less than minus two.

### Redox Homeostasis Gene Expressions

As a cell progresses toward mitotic cell cycle, cell environment becomes more oxidized [18]. MnSOD activity is high in G1/G0 phase while its activity is reduced in S/M phase. This fluctuation in MnSOD activity is inversely correlated with the level of O_2_^-^ and glucose consumption. Loss of this enzyme results in aberrant cell growth [9] because presence of superoxide accelerates cell proliferation [19]. Cyclins, CDKs and P53 interact with MnSOD and regulate its function. To check how the expression of this gene and its interacting genes change in CRC, expression results for SOD1/2/3, GPx (GPX1), Peroxiredoxin (PRRX1), Glutathione synthesis (GSS) and Glutathione reductase (GRR) were analyzed presenting in Supplementary file 1 (S1). SOD1 followed a tiny gradual increase from Normal to Primary and from Primary to Metastatic respectively (N>P&P>M) in the first dataset contrary to the second dataset, while SOD3 exhibited a gradual decrease in the first dataset from N>P&P>M contrary to the second dataset. Since, SOD1 presents in most parts of the cell especially mitochondrial intermembrane space, its tiny increase might be a defending mechanism to scavenge extra superoxide to avoid cellular damage. However, the main mitochondrial O_2_^-^ scavenger is SOD2 (MnSOD) which its expression witnesses a small reduction in primary. This would firstly demonstrate that O_2_^-^ concentration is not high in CRC. Secondly, based on what has just said, the presence of superoxide speeds up cell division. Thirdly the source of energy production in CRC cells might not come from oxidative cellular respiration. In the preprocessing step a great number of genes were removed and GSR expression remained only at the first dataset. Its expression followed a reducing trend from N>P&P>M is the first dataset while expression of GSS fluctuated with the highest value in primary. Concordance to the expression of GSS gene, GPX1 is also presenting an increase from N>P in both datasets. More reduced glutathione production by the act of GPx enzyme leads to more oxidized environment. Expressions of all these genes specifically GSR and GSS require more investigation.

CDK1 activates MnSOD and inactivates Peroxiredoxin (Prx1) by making them phosphorylated. Reduced Prx1 converts hydrogen peroxide to oxygen and water, so activity of CDK1 contributes to an increase in H_2_O_2_ production which accelerates the transition from G2 to M phase [20]. In non-irradiated dermal fibroblast cells, Cyclin B1 is present in the nucleus of G2 cells. However, following the radiation (ROS generation) Cyclin B1 and CDK1 leave the nucleus to reside in cytoplasm and mitochondria [21] and form a complex with each other which enhances the transition from G2 to M phase. CDK1 phosphorylates MnSOD to both stabilize and increase its activity. In fact, under oxidative stress in CRC, cancer cells do not overexpress SOD2, while they try to stabilize Its protein in order to remove detrimental concentrations of superoxide to balance the redox molecules pertinent for cell proliferation. In other words, Cellular redox intensity specifies whether cell moves towards apoptosis or extends its lifespan [22]. Therefore, activation and transition of CyclinB1/CDK1 complex under oxidative stress speeds up cell cycle by increasing the concentration of H_2_O_2_. What is more, under oxidative stress, P53 enters the mitochondrial matrix then phosphorylates MnSOD and reduces its antioxidant activity. This results in ROS accumulation and cell death [23]. When CDK1 phosphorylates mitochondrial P53 on serine 315, MnSOD phosphorylation by P53 is inhibited. As a result, the pertinent condition for MnSOD phosphorylation on serine 106 by CDK1 is provided which prevents apoptosis [23]. The gene expression profiles for Prx1 (PRRX1), Cyclin B1 (CCNB1) and CDK1 are presented in Supplementary file 1 (S1). Prx1 presented a total increase from N>M in the first dataset. This increase is against H_2_O_2_ increment and cell proliferation in CRC, therefore more investigation needs to be conducted on the role of Prx1 expression in CRC progression. Maybe an increase in Prx1 expression is needed to keep H_2_O_2_ at a pertinent concentration. Cyclin B1 witnessed a total increasing trend with the highest expression in the primary group in the two datasets which would augment the number of cyclin B1/CDK1 complexes. CDK1 illustrated a considerable increase from N>P&P>M in the first dataset. CDK1 expression would help to stabilize MnSOD and increase H_2_O_2_ and cyclin B1/CDK1 complex that enhance cell survival and cell proliferation. Furthermore, H_2_O_2_ bolsters the invasive behaviors of cancer cells and Epithelial to Mesenchymal Transition through overexpression of matrix metalloproteinase 2 and 9 (MMP-2 and MMP-9) and phosphorylation of ERK and NF-κB in pancreatic cancer cells [24].

Hydrogen sulfide (H_2_S) is a sulfide redox messenger and a gasotransmitter that is oxidized by dioxygenase enzymes such as sulfide quinone reductase/oxidoreductase (SQR/SQOR) and ethylmalonic encephalopathy 1 (ETHE1) [25]. Moreover, electrons are transferred from H_2_S in complexes III and IV of ETC which results in energy production. This is done by the electron carrier Coenzyme Q (Ubiquinone/CoQ) [26]. H_2_S oxidation rate depends on the interaction between enzymatic oxidation and ETC-mediated oxidation [27]. SQR enzyme with the help of CoQ uses H_2_S to oxidizes GSH to GSSH. Then, ETHE1 uses O_2_ to reduces GSSH to GSH by generating sulfite (SO3^2-^) [28, 29]. Moreover, CoQ deficiency in human skin fibroblasts results in impairment of H_2_S oxidation [26]. H_2_S is generated in human body by both cellular enzymes and anaerobic bacteria in the digestive tract [30]. That’s the reason ETHE1 and SQOR expressions are high in normal colon tissues comparing to the other tissues to detoxify the high concentration effects of hydrogen sulfide [31, 32]. The intestinal wall has a high capability of sulfide oxidation based on the activity of SQR enzyme and prevents sulfide from spreading throughout the body [30]. The expressions for ETHE1 (HSCO), SQR (SQOR) and CoQ (COQ10B) are depicted in S1. ETHE1 and SQOR pursued a similar pattern in gene expression which is the reduction from N>P&P>M. COQ10B expression followed a totally reducing trend from Normal to Metastatic (N>M). These patterns of reduction in the expression of these mitochondrial genes are so important for cancer progression. Because these enzymatic antioxidants are scavengers of H_2_S, their inhibition results in an increase in the concentration of H_2_S and one of the mechanisms of cancer cells to survive under hypoxic conditions is to produce ATP from H_2_S in the lack of oxygen [33]. Multiple studies have reported the elevated expression level of ETHE1 protein in different cancers [34]; however, our results present the opposite clues. The protein derived from the ETHE1 gene, HSCO, is a mononuclear non-heme iron persulfide dioxygenase considered as an antioxidant [35]. By binding to RelA, HSCO accelerates the translocation of this protein from nucleus to cytoplasm and thus blocks NF-κB activity [36]. Nevertheless, it is worth noting that ETHE1 exhibits anti-apoptotic activity by increasing the ubiquitination and degradation of p53 protein. In addition, HSCO, HDAC1 and p53 enzymes form a complex in which HSCO acts as a cofactor and enhances the deacetylation activity of the HDAC1 molecule on the p53 gene, thereby reducing the transcription of the p53 gene (anti-apoptotic activity of ETHE1) [34].

High concentration of hydrogen sulfide destroys complex IV in ETC [37]. However, like other redox molecules, the pivotal concentration of H_2_S is crucial for cell survival [38]. There was a minute reduction in expression of COX2 from N>P&P>M. Since COX and CoQ are electron carriers in ETC, their expression suppression triggers cells toward anaerobic respiration (Warburg Effect). Inhibition of COX2 by H_2_S [37] is another mechanism that helps cells to alter energy production route towards anaerobic respiration and lactic acid production which results in uncontrolled cell growth [39, 40]. Under specialized conditions such as cancer and oxidative stress in mammalian cells, tissue obtains its required energy from hydrogen sulfide [33]. For instance, Adenocarcinoma cells utilize sulfide for energy production [41]. H_2_S is produced from cysteine metabolism in the mitochondria using cystathionine γ-lyase (CSE) and cystathionine-β-synthase (CBS) enzymes. In fact, one of the ways to take advantage and survive from hypoxia is to produce ATP in mitochondria through H_2_S production using the CSE enzyme [38]. CSE (CSE1L) showed a considerable increase from N>P&P>M in the first dataset and it had a totally increasing trend from normal to metastatic (N>M) in the second dataset which is in favor of CRC progression. CBS expression has been shown to step up in colon and ovarian tumors [32, 42]. However, In the first dataset, CBS showed a reducing trend from N>P&P>M, while it is opposite in the second dataset. As a result, the role of CBS expression in CRC progression needs to be investigated. CSE activity depends on the translocation of this enzyme from the cytoplasm to the mitochondria by Translocase complexes located on the mitochondrial outer membrane. Translocase complex proteins transport immature proteins to one of the mitochondrial sites where they become mature proteins. Tom complex is one of the translocase complexes located on the mitochondrial outer membrane. Increased calcium leads to the transfer of the CSE enzyme from the cytosol to the mitochondria using Tom20 protein in SMC cells [38]. TOMM20 exhibited a growth from N>P&P>M in the first dataset while in the second dataset there is a total rise from N>M. Except for CBS expression, Our results for H_2_S producers CSE1L, TOM20 and H_2_S scavengers (ETHE1, SQOR) are in favor of cancer progression and suggest that sulfide molecule is produced in high concentration in CRC samples [43]. CRC cell lines treated with PAG (CSE inhibitor) and AOA (CBS inhibitor) lose viability around 60% so that these inhibitors were proposed by some researchers as a solution for inhibition of cancer progression [39, 44]. CRC cells treated with NaHS experience a rise in proliferation which is carried out through AKT and ERK phosphorylation and inhibition of tumor suppressor P21 expression [45]. Treatments of cells with NaHS for 6 hours, reduces the number of G1 cells and increases number cells in S phase [39]. in addition, Expression of lactate dehydrogenase (LDH) gene is increased proportionally to the concentration of hydrogen sulfide in IEC-6 cell lines [46]. Expression of ERK1 (MAPK3), ERK2 (MAPK1), AKT1 and p21 (CDKN1A) witnessed a reducing trend in S1. The expression results for P21 are in favor of the mentioned reports, however based on AKT and ERK expression results, the cooperation of AKT and ERK in H_2_S-mediated cell proliferation is not carried out by their expression induction in colon cancer.

NADPH oxidases in the cell membrane take one electron from NADPH and transfer it to O_2_ and produce superoxide free radicals. This family of enzymes has been shown to increase expression in many cancers [47]. Expression of NOX1/3/4 and DUOX1/2 genes in intestinal adenocarcinoma are increased [48]. Increased expression of NOX1 and DUOX2 genes in comparison with the rest of the family members are more appropriate diagnostic criteria for colon cancer [48]. NOX4 also exhibits greater expression than other genes and is highly correlated with angiogenesis, Epithelial to Mesenchymal Transition (EMT), and NOTCH signaling [48]. Expression of NOX1 exhibited a total growth from N>M with the highest value in primary in both datasets. DUOX2 experienced a noticeable growth from N>P but it is followed by a considerable reduction from Primary to Metastatic (P>M). expression induction of NOX1 and DUOX2 from N>P would indicate that, superoxide generated by these enzymes would help cancer cells to accelerate cell proliferation and metastasis. NOX4 was amongst the removed genes in the preprocessing step however, its expression was plotted separately presenting a significant increase from N>P&P>M in the first dataset. Moreover, NADPH oxidases, such as Nox1 and Nox4, are required for growth factor-mediated H_2_O_2_ production which in turn induce a number of signaling pathways, including MAP Kinase and PI3K/AKT pathways that consequently leads to cell proliferation [11]. The expression pattern of NOX1 and DUOX2 would also suggest that metastatic cells require superoxide less than primary cells.

ETHE1-overexpressed HCT116 and HT29 colon cancer cell lines show higher mitochondrial numbers than non-overexpressed cells. In addition, aerobic respiration and oxygen consumption are incremented in the overexpressed cells [49]. These effects of ETHE1 overexpression are against Warburg effect in cancer cells. In a recent study, they have shown that expression of SIRT1, PGC1α and ETHE1 is increased in FAP (familial adenomatous polyposis) intestinal samples, although the expression results in our study are totally contradictory which is based on the logic of Warburg effect. Warburg effect theory suggests that cancer cells obtain most of the ATP they need through anaerobic glycolysis no matter if they are in aerobic or in anaerobic environment [50]. Expression of SIRT1 followed a reducing trend from N>P which is followed by an increase from P>M. PGC1α (PPARGC1A) gene presented a similar pattern in expression. SIRT1 is a deacetylase that increases aerobic glycolysis [51] and PGC1α expression [52]. Therefore, SIRT1 expression reduction would prevent the induction of aerobic glycolysis which is in favor of Warburg effect. Lentiviral overexpression of ETHE1 leads to the induction of SIRT1 and PGC1α in CRC cell lines [49]. Consequently, expression induction of these three genes hinders cancer cells survival, although there should be a balance in the concentration of SIRT1 and PGC1α [49]. H_2_S augments the expression of SIRT1 in sw480 colon cancer cells [53]. Therefore, to compensate for the lack of ETHE1 activity, and keep the concentration of SIRT1 at adequate concentration, H_2_S helps to prevent the sever reduction of SIRT1 expression in both datasets.

Warburg effect is directly related to cancer development and provides a favorable environment for metastasis [54]. In glycolysis, glucose is metabolized to lactate without entering the mitochondria. Glycolysis has a faster ATP production rate but is not very effective which causes cancer cells to consume too much glucose. Thus, high glucose metabolism is one of the biomarkers of cancer cells [50]. Tumor cells produce ROS yielded from increased glycolysis which leads to pathological conditions such as cancer [55]. Although, Warburg effect is not very efficient in energy generation, as the size of tumor increases, the amount of oxygen consumed by cancer cells decreases, therefore It is an effective strategy when the tumor grows in the absence of sufficient blood vessels [50, 56, 57]. Since, mitochondrial ion and water pumps are ATP-dependent, in highly malignant tumor cells many mitochondria are eliminated as a result of decreased intracellular ATP levels. Therefore, one therapeutic approach is to increase the number of mitochondria and augment their effects on cancer cells [57].

In the nucleus, direct oxidation of Cys in the DNA-binding domain can inhibit NF-kB–DNA-binding activity. Normally, NF-kB and (inhibitor of NF-kB) IkB form a complex, which prevents the activity of NF-kB. Increased hydrogen peroxide can activate IkB-kinase (IKK) either directly through redox modification of IkK [58] or indirectly through activation of Akt and/or MEKK1, which then phosphorylates and activates IKK [59]. Activated IKK phosphorylates IKB which results in its dissociation from NF-κB. The anti-apoptotic effect of NF-κB activation is mediated by MEKK-1 (MAP3K) or Akt [60, 61]. IKB is also degraded by the ubiquitin system [62]. Because the proteasome system is also redox-sensitive, ROS can also regulate NF-κB activity by affecting the stability of IkB [63]. Expression of MAP3K followed an increase from N>P&P>M in the first dataset. AKT1 expression increased is obvious in primary in both datasets. As a result, induction of these genes in CRC would help to avoid apoptosis through NF-κB pathway.

ROS promotes the ERK1 activation and this activation increases expression of MMP2 and MMP9 [64]. Induction of MMPs proteins has been found to enhanc invasive behaviors of cancer cells [65]. Surprisingly, expression of these genes was stepped down from N>P&P>M in the first dataset. ERK1 activation also activates c-Fos which is a component of activation protein 1 (AP1). JNK is also activated by ROS which enhances c-Jun activation another component of AP1 [66]. AP1 complex enters the nucleus and activates expression of genes promoting cell proliferation [67]. The expression level for c-Fos (FOS) reduced in both datasets. c-Jun (JUN) expression fluctuates with the highest value in Primary for both datasets. In a study expression of c-Fos and c-Jun in breast tumor samples was significantly lower than adjacent non-tumor tissues [65]. The reduction of these two genes would be related to apoptotic effects of AP1 complex [66, 67]. JNK (MAPK8) showed a total reducing trend from N>M in the first dataset while p38 (MAPK14) did not show a considerable change. Based on the given information, different MAP kinase pathways would not be induced in CRC at least in gene expression level. As a result, more survey needs to be conducted on components of these pathways in CRC.

NF-κB1/2, RELA/B and REL are members of NF-κB complex. NF-κB1 and RELA presented a tiny reducing trend from N>P&P>M in the two datasets. NF-κB2 and RELB were removed in the preprocessing step and their expression showed a similar pattern. REL presented a total growth from N>M in the first dataset. inhibitor of NF-κB (NF-κBIA) also exhibits a reducing trend from N>P&P>M in the first dataset. However, Inhibitors of IKB kinase (CHUK, IKBKB) depicted an increasing trend in the first dataset. Concordance to the mentioned information on ROS-mediated NF-κB regulation, these patterns of expressions by NF-κB and its inhibitors and activators would suggest that redox-mediated regulation of NF-κB is mostly mediated post-translationally in CRC which promotes anti-apoptotic effects.

Under normal conditions, Nrf2 localizes in the cytoplasm where it interacts with the actin-binding protein Keap1. Keap1 functions as an adaptor of Cul3-based E3 ubiquitin ligase that targets Nrf2 for rapid degradation [68]. Oxidative stress is a major activator of Nrf2 pathway and dissociation of Nrf2 from Keap1 is a key step to activate of Nrf2 [68, 69]. The free Nrf2 translocates to the nucleus, heterodimerizes with Maf, and binds to a cis-acting element known as antioxidant responsive element (ARE) within the regulatory regions of many genes [70]. Nrf2 activation promotes cell survival under oxidative stress through multiple mechanisms. One major function is the transactivation of many antioxidant proteins, including peroxiredoxin 1, catalase, glutathione peroxidase, superoxide dismutase, and thioredoxin [71]. Nrf2 regulates the synthesis of glutathione. Because glutathione is not only the most abundant scavenger of ROS, but also the key controller of redox status of proteins affecting cell survival and death, the regulatory effect of Nrf2 on glutathione synthesis plays an important role in cell survival. Nrf2 (NFE2L2) did not show any significant change in the first dataset. In addition, accumulation of unfolded polypeptides after oxidative stress could also trigger apoptosis. In response to unfolded protein stress, phosphorylation of Nrf2 by PERK acts as an effector of PERK-dependent cell survival [72]. What’s more, multiple reactive cysteine residues in Keap1 are targets of modifications by ROS and electrophiles which leads to dissociation of keap1 from Nrf2 and activation of Nrf2 pathway and its translocation to the nucleus [73].

Nrf2 cannot bind to the ARE without forming a heterodimer with one of the small Maf proteins [74]; therefore, the expression level of Maf protein regulates the Nrf2–DNA-binding capacity. Interestingly, expression of Maf can also be transcriptionally regulated by Nrf2/ARE itself, thus serving as an autoregulatory feedback mechanism [75, 76]. In contrast, Bach1, a transcriptional repressor of ARE, can compete with Nrf2 to bind to the same DNA sequence, thus preventing Nrf2/ARE transcriptional activation. A recent study showed that oxidative stress can trigger nuclear accumulation of Bach1, leading to transcriptional suppression of ARE target genes [77]. Keap1 followed a totally reducing trend from N>M which would help to induce Nrf2 pathway and cell survival in CRC metastatic samples. MAF and BATCH1 exhibited a fluctuation with the lowest value in Primary for the first dataset. Expression of MAF and BATCH1 should be more surveyed.

Hypoxia-inducible factor (HIF) is an important transcription factor that regulates cellular metabolism and cell survival under hypoxic stress [2]. Under atmospheric levels of oxygen (21%), the dioxygenase PHD hydroxylates HIF-1α, which promotes the degradation of HIF-1α by proteasome system [2]. PHD needs iron in ferrous form (Fe2+) and oxygen for hydroxylation of HIF-1α. Under hypoxia, ferrous iron is converted to ferric form (Fe3+), resulting in a decrease in HIF-α hydroxylation by PHD and subsequent stabilization of HIF-α [78]. When the free HIF-1α binds to HIF-β and translocate to the nucleus, another oxygen-dependent hydroxylase enzyme called factor-inhibiting HIF-1 (FIH-1) can regulate the DNA binding and transcriptional activity of HIF. Under hypoxia, hydroxylation of HIF by FIH is hindered; therefore, HIF can be inhibited neither in cytoplasm nor in nucleus. As a result, the HIF heterodimer binds to the hypoxia-response element (HRE), and associates with coactivators such as CBP/p300. The binding results in activation or suppression of many genes involved in metabolism, angiogenesis, invasion/metastasis, and cell survival/death [79–82]. FIH-1 is able to inhibit HIF-mediated transcription of GLUT1 and VEGF-A, even under hypoxic conditions in human glioblastoma cells [83].

Interestingly, genes encoding two isoforms of PHD proteins (phd 2 & 3) are HIF targets. Therefore, HIF can also be autoregulated under hypoxia by increased expression of its regulators. This response ensures a rapid and optimal degradation of HIF-α whenever the cells return to normoxia [84]. In tumor cells, HIF plays a major role in the metabolic switch that shunts glucose metabolites from mitochondria respiration to cytosolic glycolysis (Warburg effect) [85]. HIF activation not only increases anaerobic glycolysis but also attenuates mitochondrial respiration. The former occurs through upregulation of genes encoding glucose transporter (GLUT) and lactate dehydrogenase A (LDH-A), the enzymes that convert pyruvate to lactate. The latter occurs through the induction of pyruvate dehydrogenase kinase 1 (PDK1), which inhibits pyruvate dehydrogenase, the enzyme that converts pyruvate into acetyl-CoA in the mitochondria. These two phenomena were known to prevent the entry of pyruvate to the TCA cycle and shunt pyruvate toward lactate formation through glycolysis [86, 87]. Moreover, an increase in the expression of VEGF-A demonstrates the increased rate of angiogenesis in CRC.

A study shows that silencing LDH-A resulted in a metabolic switch from glycolysis to the mitochondrial pathway and reduced tumor growth [88]. Product of LDH-A that is LDH5 is a diagnostic marker for cancer patients and is an essential target for cancer treatment as well as for increasing sensitivity to radiotherapy. LDH-A isozymes would become as future strategies for the metabolic treatment of cancer [PMID: 29913092]. These findings suggest that formation of lactate through glycolysis is important for tumor cell metabolism. Furthermore, it has been proposed that the increase in glycolysis mediated by HIF facilitates cell survival through maintaining ATP production and preventing the deleterious effect of ROS generated from mitochondrial respiration [88–90]. Under hypoxic conditions, expression of HIF1A and ARNT is supposed to be increased to enhance expression of genes involved in cancer progression although, both genes had tiny reducing trends from N>M in the first dataset. HIF-1α (HIF1A) target genes GLUT1 (SLC2A) and VEGF-A fluctuated with the highest value in primary for the first dataset while the second dataset depicted an increasing trend. inductions of these target genes would propose that under hypoxic conditions effects of HIF-1α are augmented at posttranslational level rather than mRNA level in CRC samples. PHD2 (EGLN1) exhibited a totally reducing trend from N>M in both datasets. However, PHD1 (EGLN2) was removed in the preprocessing step and it did not show any change in expression level among the groups. FIH-1 (HIF1AN) illustrated a fluctuating trend from N>M with the highest value in primary for first dataset. LDH-A was removed in the preprocessing step from both datasets. However, its expression showed a considerable increasing trend from N>P&P>M in the first dataset. This increase demonstrates that CRC cells producing ATP via anaerobic glycolysis. PDK1 followed an increasing trend from N>P&P>M in the first dataset which helps cancer cells to halt mitochondrial respiration.

## Conclusion

Despite the fact that most of the redox-related gene regulation takes place post-transcriptionally and post-translationally, in this study we tried to introduce a gene expression signature for the genes controlling redox homeostasis in CRC. To this end, we analyzed expression of 53 genes that have been reported to be affected by redox messengers. We realized that the source of energy production in CRC cells might not come from oxidative cellular respiration. Once cells divide fast, they need high energy and if all this energy is yielded by oxygen consumption, concentration of O_2_^-^ escalates which is detrimental for cancer. Moreover, concordance to this increase, cells should also soar the expression and activation of superoxide scavengers like SOD2. But, we observed that it does not happen in CRC and source of energy production must come from small molecules such as H_2_S or via cytosolic glycolysis. Totally, Expression of antioxidant enzymes such as SODs and GPx and an increase in the expression of cyclin B1/CDK1 complex helps to keep O_2_^-^ and H_2_O_2_ at an adequate concentration to create more oxidized environment suitable for cell growth in CRC tumors. Our analysis supported the presence of Warburg Effect in CRC by induction of genes that force cancer cells to move towards anaerobic glycolysis such as CSE, LDH, GLUT1 and PDK1 genes and suppression of genes that promotes aerobic respiration such as ETHE1, SQOR, Coenzyme Q and COX2. Moreover, GSR expression tended to be reduced as CRC stages progress, so rate of converting GSSG to GSH is reduced which helps to keep the oxidized environment of tumor. Furthermore, redox messengers control induction or suppression of transcription factors like NF-κB in CRC. Since NF-κB family members are reduced in expression, activation of NF-κB pathway would be controlled mostly post-translationally and it is activated by MAP kinase and AKT pathways. We also proposed that expression reduction of AP1 complex would be related to apoptotic effects of AP1. Moreover, Nrf2 pathway is more necessary in metastatic CRC tumors than primary tumors. Finally, suppression of HIF1 alpha and HIF1beta expression was observed in CRC samples and that would suggest that products of these genes might be stabilized or inhibited from degradation to play their role in promoting Warburg Effect. That’s because expression of their targets such as GLUT1, LDH and PDK1 were increased and inhibitors of HIF1A that are PHD and FIH-1 were suppressed in expression. To sum up, CRC tumors guarantee their survival and growth by creating an oxidized environment. Furthermore, under hypoxic conditions, part of the ATP generated in CRC comes from H_2_S rather than oxygen and they tend to shift cellular respiration towards anaerobic glycolysis. they produce an adequate amount of ROS which is more than in normoxia.

## References

1. Dawane, J.S. and V.A. Pandit, Understanding redox homeostasis and its role in cancer. Journal of clinical and diagnostic research: JCDR, 2012. 6(10): p. 1796.

2. Trachootham, D., et al., Redox regulation of cell survival. Antioxidants & redox signaling, 2008. 10(8): p. 1343–1374.

3. Higdon, A., et al., Cell signalling by reactive lipid species: new concepts and molecular mechanisms. Biochemical Journal, 2012. 442(3): p. 453–464.

4. Janssen-Heininger, Y.M., et al., Redox-based regulation of signal transduction: principles, pitfalls, and promises. Free Radical Biology and Medicine, 2008. 45(1): p. 1–17.

5. Sosa, V., et al., Oxidative stress and cancer: an overview. Ageing research reviews, 2013. 12(1): p. 376–390.

6. Kamata, H. and H. Hirata, Redox regulation of cellular signalling. Cellular signalling, 1999. 11(1): p. 1–14.

7. Lü, J.M., et al., Chemical and molecular mechanisms of antioxidants: experimental approaches and model systems. Journal of cellular and molecular medicine, 2010. 14(4): p. 840–860.

8. Schieber, M. and N.S. Chandel, ROS function in redox signaling and oxidative stress. Current biology, 2014. 24(10): p. R453–R462.

9. Sarsour, E.H., A.L. Kalen, and P.C. Goswami, Manganese superoxide dismutase regulates a redox cycle within the cell cycle. Antioxidants & redox signaling, 2014. 20(10): p. 1618–1627.

10. Lennicke, C., et al., Hydrogen peroxide–production, fate and role in redox signaling of tumor cells. 2015. 13(1): p. 39.

11. Sarsour, E.H., et al., Redox control of the cell cycle in health and disease. Antioxidants & redox signaling, 2009. 11(12): p. 2985–3011.

12. Kim, S.Y., et al., Regulation of singlet oxygen-induced apoptosis by cytosolic NADP+-dependent isocitrate dehydrogenase. Molecular and cellular biochemistry, 2007. 302(1-2): p. 27–34.

13. Hempel, N., P. M Carrico, and J.A. Melendez, Manganese superoxide dismutase (Sod2) and redox-control of signaling events that drive metastasis. Anti-Cancer Agents in Medicinal Chemistry (Formerly Current Medicinal Chemistry-Anti-Cancer Agents), 2011. 11(2): p. 191–201.

14. Miller, T.W., J.S. Isenberg, and D.D. Roberts, Molecular regulation of tumor angiogenesis and perfusion via redox signaling. Chemical reviews, 2009. 109(7): p. 3099–3124.

15. Irizarry, R.A., et al., Summaries of Affymetrix GeneChip probe level data. Nucleic acids research, 2003. 31(4): p. e15–e15.

16. Ramasamy, A., et al., Key issues in conducting a meta-analysis of gene expression microarray datasets. PLoS medicine, 2008. 5(9): p. e184.

17. Oldham, M.C., et al., Identification and Removal of Outlier Samples Supplement for:” Functional Organization of the Transcriptome in Human Brain. dim (dat1). 1(18631): p. 105.

18. Goswami, P.C., et al., Cell Cycle-coupled Variation in Topoisomerase IIα mRNA Is Regulated by the 3′-Untranslated Region POSSIBLE ROLE OF REDOX-SENSITIVE PROTEIN BINDING IN mRNA ACCUMULATION. Journal of Biological Chemistry, 2000. 275(49): p. 38384–38392.

19. Burdon, R.H., Superoxide and hydrogen peroxide in relation to mammalian cell proliferation. Free Radical Biology and Medicine, 1995. 18(4): p. 775–794.

20. Candas, D. and J.J. Li, MnSOD in oxidative stress response-potential regulation via mitochondrial protein influx. Antioxidants & redox signaling, 2014. 20(10): p. 1599–1617.

21. Kalen, A., et al., MnSOD and cyclin B1 coordinate a mito-checkpoint during cell cycle response to oxidative stress. Antioxidants, 2017. 6(4): p. 92.

22. Candas, D., et al., CyclinB1/Cdk1 phosphorylates mitochondrial antioxidant MnSOD in cell adaptive response to radiation stress. Journal of molecular cell biology, 2012. 5(3): p. 166–175.

23. Nantajit, D., et al., Cyclin B1/Cdk1 phosphorylation of mitochondrial p53 induces anti-apoptotic response. PloS one, 2010. 5(8): p. e12341.

24. Cao, L., et al., Curcumin inhibits H2O2-induced invasion and migration of human pancreatic cancer via suppression of the ERK/NF-κB pathway. Oncology reports, 2016. 36(4): p. 2245–2251.

25. Wróbel, M., et al., Is development of high-grade gliomas sulfur-dependent? Molecules, 2014. 19(12): p. 21350–21362.

26. Quinzii, C.M., et al., The role of sulfide oxidation impairment in the pathogenesis of primary CoQ deficiency. Frontiers in physiology, 2017. 8: p. 525.

27. Lagoutte, E., et al., Oxidation of hydrogen sulfide remains a priority in mammalian cells and causes reverse electron transfer in colonocytes. Biochimica et Biophysica Acta (BBA)- Bioenergetics, 2010. 1797(8): p. 1500–1511.

28. Mishanina, T.V., M. Libiad, and R. Banerjee, Biogenesis of reactive sulfur species for signaling by hydrogen sulfide oxidation pathways. Nature chemical biology, 2015. 11(7): p. 457.

29. Olson, K.R., Hydrogen sulfide as an oxygen sensor. Antioxidants & redox signaling, 2015. 22(5): p. 377–397.

30. Helmy, N., et al., Oxidation of hydrogen sulfide by human liver mitochondria. Nitric Oxide, 2014. 41: p. 105–112.

31. Vitvitsky, V., O. Kabil, and R. Banerjee, High turnover rates for hydrogen sulfide allow for rapid regulation of its tissue concentrations. Antioxidants & redox signaling, 2012. 17(1): p. 22–31.

32. Szabo, C., et al., Tumor-derived hydrogen sulfide, produced by cystathionine-β-synthase, stimulates bioenergetics, cell proliferation, and angiogenesis in colon cancer. Proceedings of the National Academy of Sciences, 2013. 110(30): p. 12474–12479.

33. Szabo, C., et al., Regulation of mitochondrial bioenergetic function by hydrogen sulfide. Part I. Biochemical and physiological mechanisms. British journal of pharmacology, 2014. 171(8): p. 2099–2122.

34. Higashitsuji, H., et al., Enhanced deacetylation of p53 by the anti-apoptotic protein HSCO in association with histone deacetylase 1. Journal of Biological Chemistry, 2007. 282(18): p. 13716–13725.

35. Salomone-Stagni, M., Biochemical and Biophysical Characterization of the Human Ethylmalonic Encephalopathy non-Heme Sulfur [Fe]-Dioxygenase ETHE1, and X-ray Absorption Spectroscopy Applications and Methods Development. 2010.

36. Higashitsuji, H., et al., A novel protein overexpressed in hepatoma accelerates export of NF-κB from the nucleus and inhibits p53-dependent apoptosis. Cancer cell, 2002. 2(4): p. 335–346.

37. Cooper, C.E. and G.C. Brown, The inhibition of mitochondrial cytochrome oxidase by the gases carbon monoxide, nitric oxide, hydrogen cyanide and hydrogen sulfide: chemical mechanism and physiological significance. Journal of bioenergetics and biomembranes, 2008. 40(5): p. 533.

38. Fu, M., et al., Hydrogen sulfide (H2S) metabolism in mitochondria and its regulatory role in energy production. Proceedings of the National Academy of Sciences, 2012. 109(8): p. 2943–2948.

39. Wang, C.-N., et al., CBS and CSE are critical for maintenance of mitochondrial function and glucocorticoid production in adrenal cortex. Antioxidants & redox signaling, 2014. 21(16): p. 2192–2207.

40. Annane, D., et al., A 3-level prognostic classification in septic shock based on cortisol levels and cortisol response to corticotropin. Jama, 2000. 283(8): p. 1038–1045.

41. Szczesny, B., et al., Hydrogen sulfide in lung adenocarcinoma: a tumor cell survival factor, an enhancer of mitochondrial DNA repair and a cellular bioenergetic stimulator. Nitric Oxide, 2015(47): p. S46.

42. Renga, B., Hydrogen sulfide generation in mammals: the molecular biology of cystathionine-β-synthase (CBS) and cystathionine-γ-lyase (CSE). Inflammation & Allergy-Drug Targets (Formerly Current Drug Targets-Inflammation & Allergy), 2011. 10(2): p. 85–91.

43. Caliendo, G., et al., Synthesis and biological effects of hydrogen sulfide (H2S): development of H2S-releasing drugs as pharmaceuticals. Journal of medicinal chemistry, 2010. 53(17): p. 6275–6286.

44. Rodrigues, C. and S.S. Percival, Immunomodulatory effects of glutathione, garlic derivatives, and hydrogen sulfide. Nutrients, 2019. 11(2): p. 295.

45. Murugan, R.S., et al., Intrinsic apoptosis and NF-κB signaling are potential molecular targets for chemoprevention by black tea polyphenols in HepG2 cells in vitro and in a rat hepatocarcinogenesis model in vivo. Food and Chemical Toxicology, 2010. 48(11): p. 3281–3287.

46. Wu, D., et al., Hydrogen sulfide in cancer: friend or foe? Nitric oxide, 2015. 50: p. 38–45.

47. Roy, K., et al., NADPH oxidases and cancer. Clinical Science, 2015. 128(12): p. 863–875.

48. Cho, S.Y., et al., Expression of NOX Family Genes and Their Clinical Significance in Colorectal Cancer. Digestive diseases and sciences, 2018. 63(9): p. 2332–2340.

49. Witherspoon, M., et al., ETHE1 overexpression promotes SIRT1 and PGC1α mediated aerobic glycolysis, oxidative phosphorylation, mitochondrial biogenesis and colorectal cancer. Oncotarget, 2019. 10(40): p. 4004–4017.

50. Kobayashi, Y., et al., Warburg effect in Gynecologic cancers. Journal of Obstetrics and Gynaecology Research, 2019. 45(3): p. 542–548.

51. Li, X., SIRT1 and energy metabolism. Acta Biochim Biophys Sin, 2013. 45(1): p. 51–60.

52. Fernandez-Marcos, P.J. and J. Auwerx, Regulation of PGC-1α, a nodal regulator of mitochondrial biogenesis. The American journal of clinical nutrition, 2011. 93(4): p. 884S–890S.

53. Kumarasamy, A. and G. Kurian, Hydrogen Sulfide Promotes Proliferation of HT-29 Colon Cancer Cells in a Mitochondria-independent Pathway. Indian J Pharm Sci, 2019. 81(3): p. 456–463.

54. Lu, J., M. Tan, and Q. Cai, The Warburg effect in tumor progression: mitochondrial oxidative metabolism as an anti-metastasis mechanism. Cancer letters, 2015. 356(2): p. 156–164.

55. Lebelo, M.T., A.M. Joubert, and M.H. Visagie, Warburg effect and its role in tumourigenesis. Archives of pharmacal research, 2019: p. 1–15.

56. Ganapathy-Kanniappan, S. and J.-F.H. Geschwind, Tumor glycolysis as a target for cancer therapy: progress and prospects. Molecular cancer, 2013. 12(1): p. 152.

57. Schwartz, L., C. T Supuran, and K. O Alfarouk, The Warburg effect and the hallmarks of cancer. Anti-Cancer Agents in Medicinal Chemistry (Formerly Current Medicinal Chemistry-Anti-Cancer Agents), 2017. 17(2): p. 164–170.

58. Kamata, H., et al., Hydrogen peroxide activates IκB kinases through phosphorylation of serine residues in the activation loops. FEBS letters, 2002. 519(1-3): p. 231–237.

59. Madrid, L.V., et al., Akt stimulates the transactivation potential of the RelA/p65 subunit of NF-κB through utilization of the IκB kinase and activation of the mitogen-activated protein kinase p38. Journal of Biological Chemistry, 2001. 276(22): p. 18934–18940.

60. Nawata, R., et al., MEK kinase 1 mediates the antiapoptotic effect of the Bcr-Abl oncogene through NF-κB activation. Oncogene, 2003. 22(49): p. 7774.

61. Vandermoere, F., et al., The antiapoptotic effect of fibroblast growth factor-2 is mediated through nuclear factor-κB activation induced via interaction between Akt and IκB kinase-β in breast cancer cells. Oncogene, 2005. 24(35): p. 5482.

62. Serasanambati, M. and S.R. Chilakapati, Function of nuclear factor kappa B (NF-kB) in human diseases-a review. South Indian Journal of Biological Sciences, 2016. 2(4): p. 368–387.

63. D’Ignazio, L. and S. Rocha, Hypoxia induced NF-κB. Cells, 2016. 5(1): p. 10.

64. Bautista-López, N., et al., Induction of Increased Levels of Matrix Metalloproteinase-2 (MMP-2) and 9 in Human Breast Cancer Cell Lines by Activation of GM-CSF Receptor Bc via C-Fos–ERK 1/2 Signaling. J Clin Exp Oncol 6, 2017. 2: p. 2.

65. Li, Z., et al., Activation of MMP-9 by membrane type-1 MMP/MMP-2 axis stimulates tumor metastasis. Cancer science, 2017. 108(3): p. 347–353.

66. Weekes, D., et al., Regulation of osteosarcoma cell lung metastasis by the c-Fos/AP-1 target FGFR1. Oncogene, 2016. 35(22): p. 2852.

67. Chatterjee, K., et al., EGFR-mediated matrix metalloproteinase-7 up-regulation promotes epithelial–mesenchymal transition via ERK1-AP1 axis during ovarian endometriosis progression. The FASEB Journal, 2018. 32(8): p. 4560–4572.

68. Niture, S.K., et al., Nrf2 signaling and cell survival. Toxicology and applied pharmacology, 2010. 244(1): p. 37–42.

69. Ma, Q., Role of nrf2 in oxidative stress and toxicity. Annual review of pharmacology and toxicology, 2013. 53: p. 401–426.

70. Jaramillo, M.C. and D.D. Zhang, The emerging role of the Nrf2–Keap1 signaling pathway in cancer. Genes & development, 2013. 27(20): p. 2179–2191.

71. Ishii, T., et al., Transcription factor Nrf2 coordinately regulates a group of oxidative stress-inducible genes in macrophages. Journal of Biological Chemistry, 2000. 275(21): p. 16023–16029.

72. Bobrovnikova-Marjon, E., et al., PERK promotes cancer cell proliferation and tumor growth by limiting oxidative DNA damage. Oncogene, 2010. 29(27): p. 3881.

73. Huang, H.-C., T. Nguyen, and C.B. Pickett, Phosphorylation of Nrf2 at Ser-40 by protein kinase C regulates antioxidant response element-mediated transcription. Journal of Biological Chemistry, 2002. 277(45): p. 42769–42774.

74. Motohashi, H., et al., Small Maf proteins serve as transcriptional cofactors for keratinocyte differentiation in the Keap1–Nrf2 regulatory pathway. Proceedings of the National Academy of Sciences, 2004. 101(17): p. 6379–6384.

75. Canning, P., F.J. Sorrell, and A.N. Bullock, Structural basis of Keap1 interactions with Nrf2. Free Radical Biology and Medicine, 2015. 88: p. 101–107.

76. Katsuoka, F., et al., Genetic evidence that small maf proteins are essential for the activation of antioxidant response element-dependent genes. Molecular and cellular biology, 2005. 25(18): p. 8044–8051.

77. Muto, A., et al., Activation of Maf/AP-1 repressor Bach2 by oxidative stress promotes apoptosis and its interaction with promyelocytic leukemia nuclear bodies. Journal of Biological Chemistry, 2002. 277(23): p. 20724–20733.

78. Bell, E.L. and N.S. Chandel, Mitochondrial oxygen sensing: regulation of hypoxia-inducible factor by mitochondrial generated reactive oxygen species. Essays in biochemistry, 2007. 43: p. 17–28.

79. Zhao, T., et al., LASP1 is a HIF1α target gene critical for metastasis of pancreatic cancer. Cancer research, 2015. 75(1): p. 111–119.

80. Du, R., et al., HIF1α induces the recruitment of bone marrow-derived vascular modulatory cells to regulate tumor angiogenesis and invasion. Cancer cell, 2008. 13(3): p. 206–220.

81. Kim, J.Y., J.-K. Kim, and H. Kim, ABCB7 simultaneously regulates apoptotic and non-apoptotic cell death by modulating mitochondrial ROS and HIF1α-driven NFκB signaling. Oncogene, 2019: p. 1–14.

82. Han, J.E., et al., Inhibition of HIF1α and PDK Induces Cell Death of Glioblastoma Multiforme. Experimental neurobiology, 2017. 26(5): p. 295–306.

83. Wang, E., et al., The role of factor inhibiting HIF (FIH-1) in inhibiting HIF-1 transcriptional activity in glioblastoma multiforme. PLoS One, 2014. 9(1): p. e86102.

84. Brahimi-Horn, M.C. and J. Pouysségur, Hypoxia in cancer cell metabolism and pH regulation. Essays in biochemistry, 2007. 43: p. 165–178.

85. Lu, H., R.A. Forbes, and A. Verma, Hypoxia-inducible factor 1 activation by aerobic glycolysis implicates the Warburg effect in carcinogenesis. Journal of Biological Chemistry, 2002. 277(26): p. 23111–23115.

86. Brahimi-Horn, M.C. and J. Pouysségur, Oxygen, a source of life and stress. FEBS letters, 2007. 581(19): p. 3582–3591.

87. Kim, J.-w., et al., HIF-1-mediated expression of pyruvate dehydrogenase kinase: a metabolic switch required for cellular adaptation to hypoxia. Cell metabolism, 2006. 3(3): p. 177–185.

88. Ooi, A.T. and B.N. Gomperts, Molecular pathways: targeting cellular energy metabolism in cancer via inhibition of SLC2A1 and LDHA. Clinical Cancer Research, 2015. 21(11): p. 2440–2444.

89. Batra, S., et al., Cancer metabolism as a therapeutic target. Oncology, 2013. 27(5).

90. Zheng, J., Energy metabolism of cancer: Glycolysis versus oxidative phosphorylation. Oncology letters, 2012. 4(6): p. 1151–1157.

